# PyMOL plugin for Protein Circuit Topology

**DOI:** 10.1101/2025.10.21.683762

**Authors:** Matīss Dimiņš, Alexander Bazba, Ádám Mogyorósi, Ella Kennon, Tomás Díaz Fiol, Leïla Aïkili Hagen, Vahid Sheikhhassani, Vasily Akulov, Alireza Mashaghi

## Abstract

Circuit Topology (CT) provides a fundamental framework for analysing folded polymer chains, with applications in functional annotation, protein engineering and drug development. We present a protein CT analysis plugin for PyMOL v3.1.6.1 with a graphical user interface (GUI), automatic installation, and novel features developed through integration with PyMOL’s application programming interface (API). The plugin integrates various previously developed CT methodologies for studying structured proteins and their complexes as well as the dynamics of disordered proteins. Analysis of a representative protein and a molecular dynamics trajectory demonstrates the plugin’s three analysis modes and their outputs. The plugin reproduces the reference ProteinCT implementation exactly on the structures tested, and is distributed with a versioned release, a pinned environment and a one-command reproduction of every static result reported here.

## 1. Motivation and significance

Circuit topology is a mathematical and conceptual framework used to formally describe the topology of protein molecules [1]. The framework forms the basis of a general classification and analysis approach that can be used to analyse and study the structures of folded linear chains, including structured proteins, intrinsically disordered proteins, and beyond [2-5]. Computational implementations of Circuit Topology have enabled researchers to apply this framework to the analysis of protein structures, facilitating quantitative investigations of topological organization and structural complexity [6].

Given the wide usage of Circuit Topology and despite powerful computational developments, there is a need for accessible and integrated software for its application to protein analysis. Researchers without extensive technical familiarity with Python, GitHub, or Anaconda may find computational workflows challenging to adopt, while topology analysis and molecular visualization often require the use of separate platforms such as PyMOL or VMD. An integrated, user-friendly environment that combines Circuit Topology analysis with molecular visualization would therefore be highly beneficial.

Here, we present a Python-based plugin for the molecular visualization platform PyMOL that enables Circuit Topology analysis directly within a widely used molecular visualization environment. The plugin provides an intuitive graphical user interface (GUI) that streamlines the execution of Circuit Topology workflows while maintaining compatibility with established protein structure analysis practices. The plugin incorporates various established CT tools [6] as well as a recently developed CT Folding Score proposed by Akulov et al., enabling compaction analysis of intrinsically disordered protein chains [7]. As such, this report introduces our plugin as a powerful software for performing protein Circuit Topology.

## 2. Software Description

### 2.1 Software Architecture

The plugin is organized into distinct modules for calculation, plotting, and exporting (Fig. 1 (left)), allowing the same backend routines to be called from the three GUI tabs (Fig. 2). The three GUI tabs correspond to the three CT analysis modes: Local Circuit Topology (LCT), Single-File Analysis, and Multi-File Analysis. Each analysis mode is connected to a backend that parses and prepares proteins (polymer atoms), detects contact pairs, performs (circuit) topology classification, generates plots, and exports tables. Input structures can be provided either from objects already imported into PyMOL or from external structure files (PDB/CIF) imported in the GUI. Analysis is performed on the polymer atoms of the selected object: waters, ions, ligands and other non-polymer atoms are excluded automatically and reported to the user, and the user’s structure is never modified. Contact pairs are detected between heavy atoms (hydrogen atoms are ignored by default), using a 4.5 Å distance criterion, a minimum of five atom-atom contacts per residue pair, and exclusion of residues separated by three positions or fewer along the chain; all three are adjustable by the user. An optional “Count hydrogen atoms (ProteinCT-identical)” setting counts every atom present, which reproduces the reference ProteinCT implementation on hydrogen-bearing files at the cost of comparability between models with and without hydrogens. A full methodological specification, including the handling of hydrogens, alternate conformers, insertion codes, unresolved loops and chemically modified residues, is distributed with the software.

**Fig. 1:**
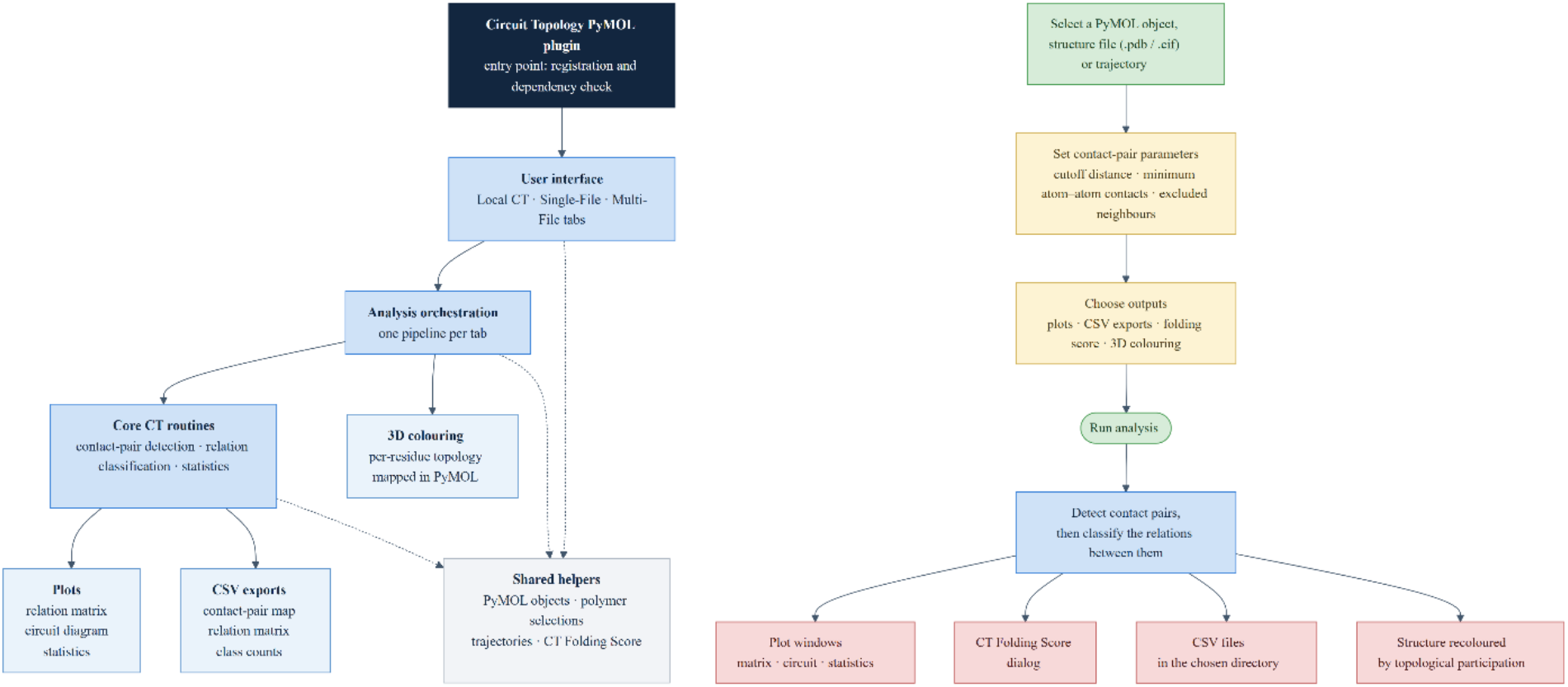
Plugin architecture (left) and analysis workflow (right). The plugin architecture shows the modules of the plugin and the relations between them. Each module groups several source files by responsibility. The analysis workflow details the steps a user follows in performing a CT analysis, tracing inputs to outputs and showing the decisions required at each step.

**Fig. 2:**
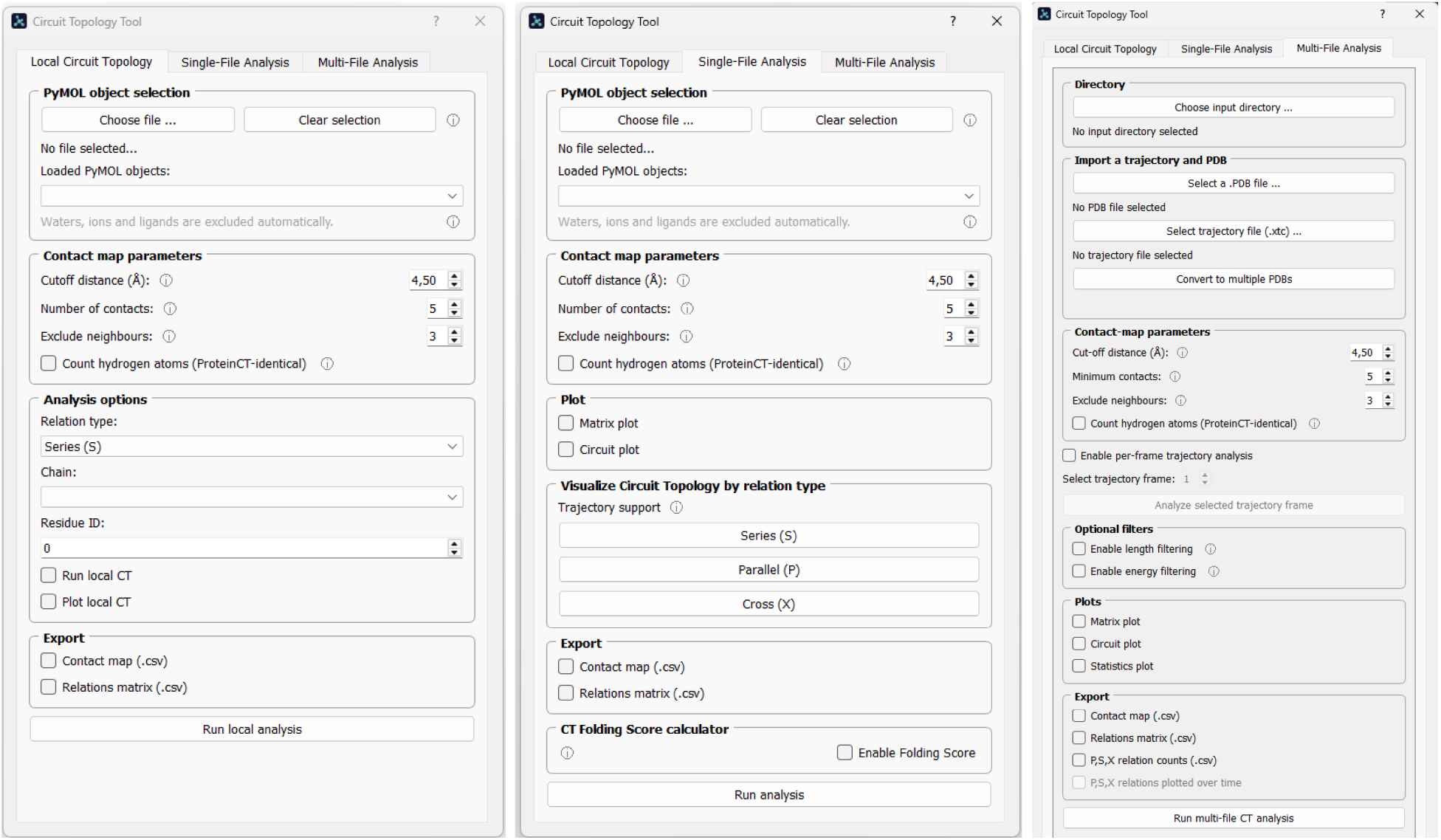
The three Circuit Topology plugin analysis tabs. From left to right, “Local Circuit Topology”, “Single-File Analysis”, and “Multi-File Analysis” tabs.

At the core of the analysis is a contact pair map derived from a geometric criterion (distance threshold) combined with simple filters that suppress trivial contact pairs. For a single chain, the resulting set of intrachain contact pairs is converted into CT information by classifying the relation between every two contact pairs into the standard relation types (Series (S), Parallel (P) and its inverse, Cross (X), and their concerted variants), which depend on the ordering of the contact pair endpoints along the sequence rather than on global orientation [6]. For an object with several chains, the plugin analyses the whole model at once: the neighbour search runs over all chains, so inter-chain contact pairs are included, and the residues of all chains are placed on one concatenated index (the neighbour-exclusion window therefore also spans a chain boundary). A relation is assigned S, P or X only when both of its contact pairs lie within one and the same chain. Every other relation, that is, every relation in which at least one of the two contact pairs joins two chains, is classified with the multi-chain vocabulary of Heidari et al. [8], as adapted in ProteinCT [6]: Independent (I) when the two contact pairs share no chain, Tandem (T) when they share a chain but cannot be joined into a closed path along chains and contacts, and Loop (L) when they can, which for two contact pairs means both join the same two chains. The digit in the exported labels (I2, I3, I4, T2, T3) is the number of chains the two contact pairs span, the superscript of Heidari et al.’s notation. The two vocabularies are not interchangeable, and the level is chosen automatically from the number of chains. Per-residue colouring, the CT Folding Score, Local Circuit Topology, length and energy filtering, and the contact-pair CSV export are computed for each chain independently from its intrachain contact pairs. This separation between contact pair detection and topology classification keeps the workflow transparent: users can tune how “contact pairs” are defined and immediately see how the topological summary changes. Fig. 1 (right) collapses the three analysis modes into one abstracted CT analysis workflow, detailing the decisions a user is required to make at each step of the analysis from input to output.

### 2.2 Software Functionalities

The primary purpose of this PyMOL plugin is to offer a user-friendly integration of the methods described in the MethodsX paper “ProteinCT: An implementation of the protein circuit topology framework” [6], as well as to include additional CT functionalities developed since its publication. The ProteinCT toolkit required the use of Anaconda, GitHub and JupyterLabs, which limited its accessibility. The plugin carries over the ProteinCT base analysis unchanged: contact-pair map construction (with hydrogen atoms ignored by default, Section 2.1), classification of contact-pair relations, the fraction of P + X relations that ProteinCT reports as “entanglement statistics”, the circuit diagram, the relation heat map —all reproduce the reference implementation on the hydrogen-free structures tested. The optional all-atom mode reproduces the original ProteinCT analysis on hydrogen-bearing structures as well. A parity example is supplied in Supplementary Fig. S1. We verify this automatically on every commit (Section 2.3). Four capability groups of the reference toolkit are not carried over in this release: secondary-structure contact pair maps and their filters, which depend on external secondary-structure assignment programs; contact pair order and the globularity score; alternative contact pair definitions based on residue centres of mass or geometry; and the file-retrieval and structure-preparation helpers, whose function is provided natively by PyMOL. Everything else in the plugin is new relative to the reference toolkit: the graphical interface and its three analysis modes, ability to operate directly on structures already loaded in the viewer, automatic dependency installation, non-destructive input handling, trajectory and batch analysis, the projection of per-residue topology onto the 3D structure with a labelled scale, and the CT Folding score of Akulov et al. [7]. Table 1 summarises the division.

**Table 1:** Comparison of capabilities between the original ProteinCT method and the new ProteinCT PyMOL plugin.

| Capability | ProteinCT | This plugin | Status |
| --- | --- | --- | --- |
| Contact-pair map (single chain and whole model) | Yes | Yes | **Inherited.** Hydrogen atoms are ignored by default. Optionally, users can toggle ProteinCT-identical all-atom mode. |
| Contact-pair relation matrix (S, P, $P^{-1}$, X, CP, $CP^{-1}$, CS) | Yes | Yes | **Inherited, identical** |
| Fraction of P + X relations vs. distance from diagonal (ProteinCT's "entanglement statistics") | Yes | Yes | **Inherited, identical** |
| Circuit (arc) diagram | Yes | Yes | **Inherited, identical** |
| Relation-matrix heat map | Yes | Yes | **Inherited and corrected.** Fixed color scale and each relation type now has its own colour. |
| Local circuit topology | Yes | Yes | **Inherited.** Dedicated tab with chain and residue pickers. |
| Contact-pair length filtering | Yes | Yes | **Inherited** |
| Contact-pair energy filtering | Yes | Yes | **Inherited** |
| CSV export of contact pair map, relation matrix and relation-type count list | Yes | Yes | **Inherited, corrected.** Cross is consistently exported as 'X' now. |
| Secondary-structure contact pair maps and filtering (STRIDE / DSSP) | Yes | No | **Not carried over.** Requires externally downloaded STRIDE files and a DSSP binary. |
| Contact pair order; globularity score; centre-of-mass and centre-of-geometry contact pair maps; circuit export | Yes | No | **Not carried over in this release.** Not essential to the ProteinCT method. |
| mmCIF download, batch download, PDB pre-preparation, static PyMOL-script generation | Yes | No | **Redundant.** PyMOL performs these natively. |
| Graphical user interface | No | Yes | **New** |
| Runs inside a molecular viewer, on objects already loaded | No | Yes | **New** |
| Automatic dependency installation | No | Yes | **New** |
| Non-destructive input handling | No | Yes | **New** |
| Trajectory support: load, analyse any frame, analyse all frames, optional frame export | No | Yes | **New** |
| Batch analysis over a folder of structures | No | Yes | **New** |
| Per-residue topology mapped onto the 3D structure, with a labelled colour scale | No | Yes | **New** |
| CT Folding Score | No | Yes | **New** – implements the published metric of Akulov et al. [7] |
| Automated test suite, cross-platform installation testing, reproducible capsule | No | Yes | **New** |

### 2.3 Validation, installation testing and reproducibility

Installation is verified automatically for each release. On Windows, Linux and macOS a clean machine (via GitHub Actions) installs PyMOL 3.1.6.1 from the official installer. The plugin itself is installed through PyMOL’s own Plugin Manager, and the full test suite is executed inside that installation. The platforms automatically tested are Windows Server 2025, Ubuntu 24.04 and macOS 15. Installation and interactive use via the GUI were additionally verified manually on Windows 10, Windows 11, Ubuntu Linux 24.04, and macOS 15. On Apple Silicon, PyMOL is itself an x86-64 application running under Rosetta 2, since no native arm64 build is published. The plugin installs only the packages missing from PyMOL’s own bundled conda environment. The plugin resolves the required missing package versions by pinning Python and PyMOL versions, thus ensuring no changes to the existing dependencies in the environment. If PyMOL was installed system-wide on the device, which the plugin detects in advance, then the plugin requires elevated privileges. As an alternative, a manual installation route is documented.

The plugin divides the ProteinCT framework into three tabs according to three different analysis modes. The LCT tab (Fig. 2, left) adapts local circuit topology analysis, performing ProteinCT on individual residues [6]. The Single-File Analysis tab (Fig. 2, middle) is the standard ProteinCT analysis mode, performing CT analysis on an entire object. It has been extended with the CT Folding Score [7] and additional visualization options. Both the local and single-file analysis tabs accept PDB and CIF files uploaded by the user along with objects already loaded into PyMOL. The Multi-File Analysis tab (Fig. 2, right) accepts two input modes: a folder of PDB/CIF files or a trajectory loaded into PyMOL alongside its structure file. A trajectory is analysed directly from its frames, and no conversion to individual structure files is required. Exporting frames as separate PDB files is available for users who want them on disk. Individual frames can also be analysed from the ‘Single-File Analysis’ or ‘Local Circuit Topology’ tabs by selecting the trajectory object and using PyMOL’s built-in frame controls. Each tab has several customizable parameters which can be seen in Fig. 2. The user can also refer to the plugin manual for a more in-depth guide with additional instructions on installation and usage.

## 3. Illustrative Examples

The Circuit Topology plugin offers three analysis modes: LCT examines topological relationships around individual residues to identify folding motifs, Single-File Analysis characterizes the complete topology of a protein structure for comparative studies, and Multi-File Analysis tracks topological changes across trajectories to monitor conformational transitions. Below we demonstrate the two static analysis modes using hen egg-white lysozyme (PDB 1AKI), a well-resolved, experimentally determined single-chain globular protein without deposited hydrogen atoms (Figs. 3-6), and the Multi-file analysis time-resolved output using a molecular dynamics trajectory of the intrinsically disordered AF1c fragment of the glucocorticoid receptor (Fig. 7). The AF1c simulation protocol (GROMACS, amber99SBdisp force field, 150 mM NaCl, 310 K, 1 µs per replica) is described in the study from which the trajectory is taken [7]. The frames analysed here are the 205 frames of replica 3 of the 150 mM NaCl simulation of the S203P phosphovariant, taken from the archive 150mM_S203P.7z of the deposited dataset [9]. The trajectory file includes hydrogens.

**Fig. 3:**
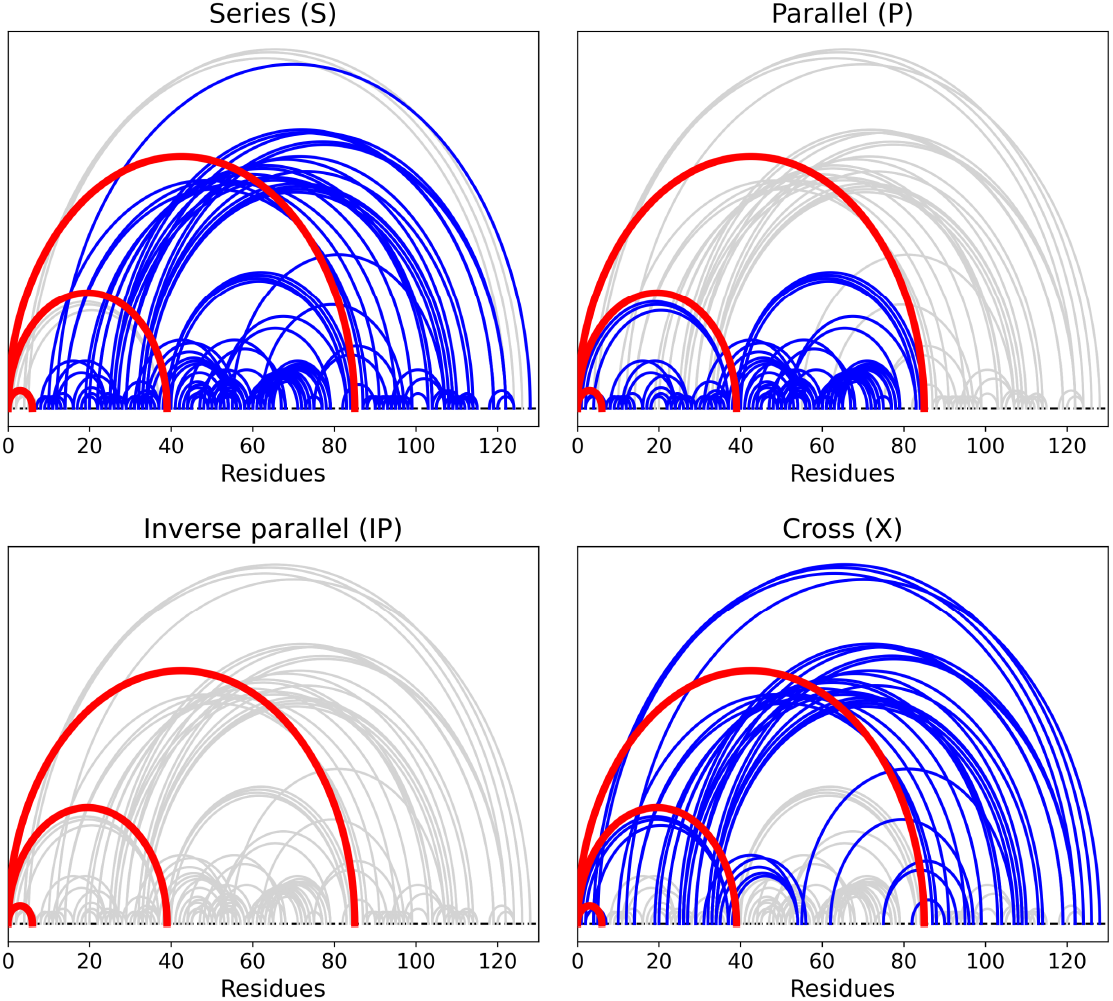
The four CT relation types (S, P, IP / P^−1^, X) visualized in circuit plots for hen egg-white lysozyme (PDB 1AKI, chain ‘A’, residue ID: 1). In each panel the contact pairs having the target residue as an endpoint are drawn in red, the contact pairs standing in the named relation to them are drawn in blue, and all remaining contact pairs of the structure are drawn in grey. The relation between two contact pairs is determined solely by the order of their four endpoints along the sequence: nested endpoints give Parallel and its inverse, disjoint endpoints give Series, and interleaved endpoints give Cross.

### 3.1 Local Circuit Topology: Relation Types Around a Residue

LCT analysis enables targeted investigation of topological organization around individual residues, which is particularly valuable for identifying structural motifs critical to protein folding or function. To demonstrate this capability, we sampled an example residue from hen egg-white lysozyme (PDB 1AKI, chain ‘A’, residue ID: 1). 1AKI is an X-ray structure without deposited hydrogen atoms, therefore the hydrogen setting does not affect any lysozyme result in this section and Section 3.2.

The structure was loaded into PyMOL and analyzed using the LCT tab. We set the cutoff distance to 4.5 Å (Angstroms), which is standard for detecting residues in proximity that form stable structural contacts. This value avoids counting residues that are merely nearby by chance. The minimum of five atomic contacts per residue contact pair ensures that only meaningful interactions are included, filtering out weak or accidental contacts. We excluded three sequential neighbours because consecutive residues along the protein backbone are always close together regardless of folding—including them would add many uninformative contacts that obscure the interesting structural features. These default parameters work well for most proteins; however, users can modify them as needed. For example, when analysing flexible proteins or investigating weak contacts, users may increase the cutoff distance to capture weaker, more transient interactions.

At this point, we specify the chain for this PDB object (‘A’) and select the residue with ID: 1. The choice of residue is trivial and is done for illustrative purposes. We enabled both ‘Run local CT’ and ‘Plot local CT’ options and executed the analysis with the ‘Run local analysis’ button. We did this for all relation types offered by LCT: Series (S), Parallel (P), Inverse Parallel (IP / P^−1^), Cross (X). The plugin computed the topological classification of all contact pairs involving the selected residue and generated visualization plots.

The resulting visualization (Fig. 3) shows four circuit plots comparing relation types at the same residue. For the circuit plot visualizing relation type ‘Parallel (P)’, the residue contact pairs that either end or begin at the target residue are curves highlighted in red. In the same plot, the residue contact pairs highlighted as blue curves are contact pairs that are in a Parallel relation with at least one contact pair highlighted in red, meaning that their endpoints are nested within one another. In contrast, the circuit plot for the ‘Cross (X)’ relation highlights, in blue, every contact pair whose endpoints interleave in sequence with those of the target contact pairs, shown in red. A Cross relation is a statement about the ordering of contact endpoints along the sequence: the two contact pairs can be neither nested within one another nor placed one after the other. It does not by itself imply a physical crossing of the chain, a threading event, or a knot. Establishing any of those would require a geometric entanglement measure, which circuit topology does not compute. The circuit plot for the ‘Series (S)’ relation shows the contact pairs that lie entirely before or after the target contact pairs along the sequence, whereas the plot for ‘Inverse Parallel (IP)’ shows the evident inverse of the ‘Parallel (P)’ relation - it highlights the contact pairs (in blue) inside which the target contact pairs (in red) are nested. Evidently, for this residue there are none. For all circuit plots, the greyed-out curves are all the other contact pairs of the structure that did not form the selected relation with the target residue.

### 3.2 Single-File Analysis: Comprehensive Topology Characterization

Single-file analysis characterizes the circuit topology of an entire protein structure, producing contact-pair matrices, circuit diagrams, and a quantitative folding score. We applied this mode to analyze the complete lysozyme structure.

After loading the structure, we applied the same contact-map parameters as in the LCT analysis (4.5 Å cutoff distance, minimum of five contacts, and exclusion of three sequential neighbours), since these values are appropriate defaults for detecting stable contact pairs in folded proteins. We selected both ‘Matrix Plot’ and ‘Circuit Plot’ visualization options, enabled the CT Folding Score calculation by checking ‘Enable Folding Score’, and selected both export options. Upon running the analysis, the plugin generated two visualizations (Fig. 4): a color-coded relation matrix showing the type of every contact pair relation across the protein sequence, and a circuit plot representing these contact pairs as curves connecting residue pairs.

**Fig. 4:**
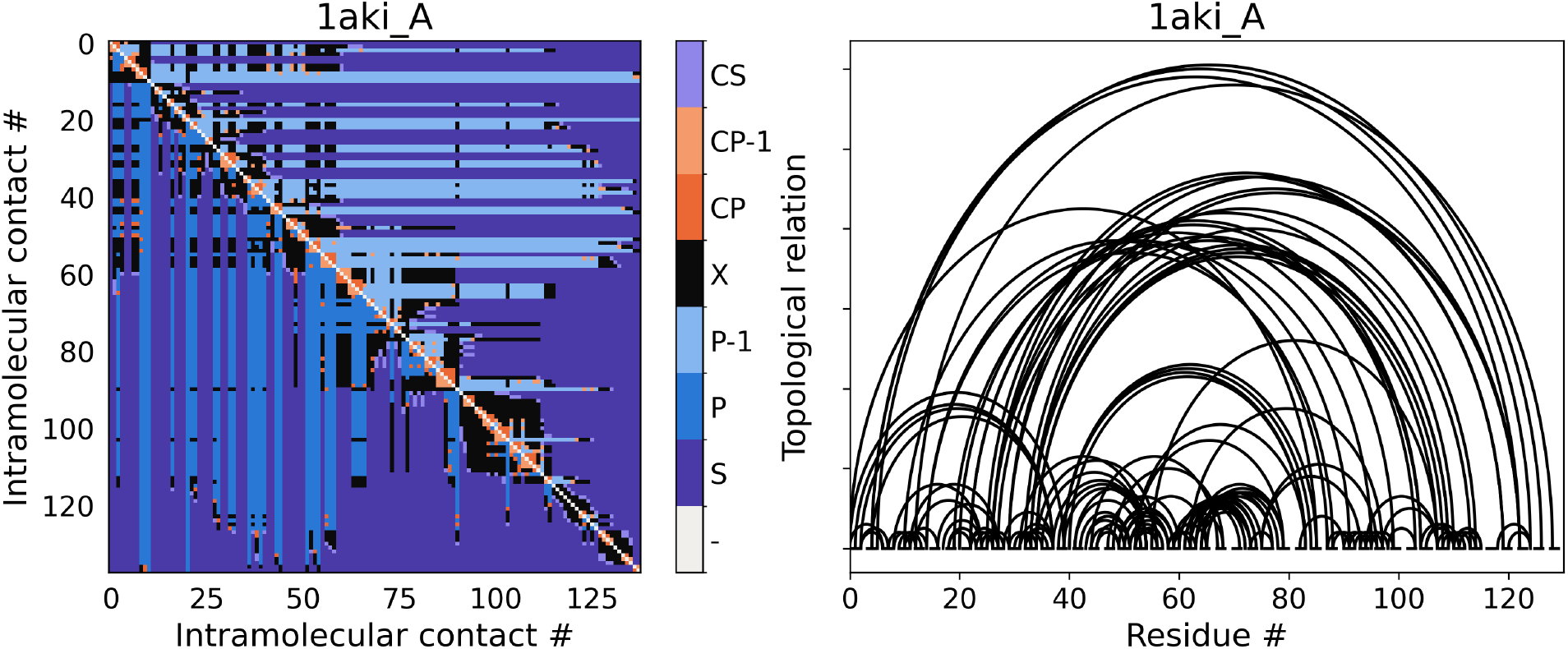
Relation matrix (left) and circuit plot (right) generated during the single-file analysis of hen egg-white lysozyme (PDB 1AKI, chain A), providing graphical summaries of the topological relations between contact pairs. Each of the eight relation types in the matrix has its own colour. The colour assigned to a relation type does not depend on which types the current structure contains.

The relation matrix showed the distribution of contact-pair relation types across the structure, and the circuit plot showed all contact pairs as curves. To map topology onto the three-dimensional structure, we activated the ‘Visualize Circuit Topology by relation type’ feature. The plugin coloured residues according to their level of participation in the Series (S), Parallel (P) and Cross (X) relations (Fig. 5). Residues with no participation in the selected relation type are white, and non-zero values are assigned to five graded levels. The colour bar placed in the viewer is labelled with the minimum and maximum of the scale, and the level bounds are printed to the PyMOL log and given in the caption of Fig. 5. Because each map is scaled to its own distribution, colours are comparable within a panel but not between panels. Colouring is applied to a copy of the object, so the original structure keeps its deposited B factors and can be restored by deleting the copy. Results for the lysozyme single-file CT analysis were exported as CSV files containing the relation matrix and the contact-pair map (see Supplementary Fig. S2 for the relation matrix). Finally, the CT Folding Score is both printed to standard output in PyMOL and shown as a pop-up GUI dialog.

**Fig. 5:**
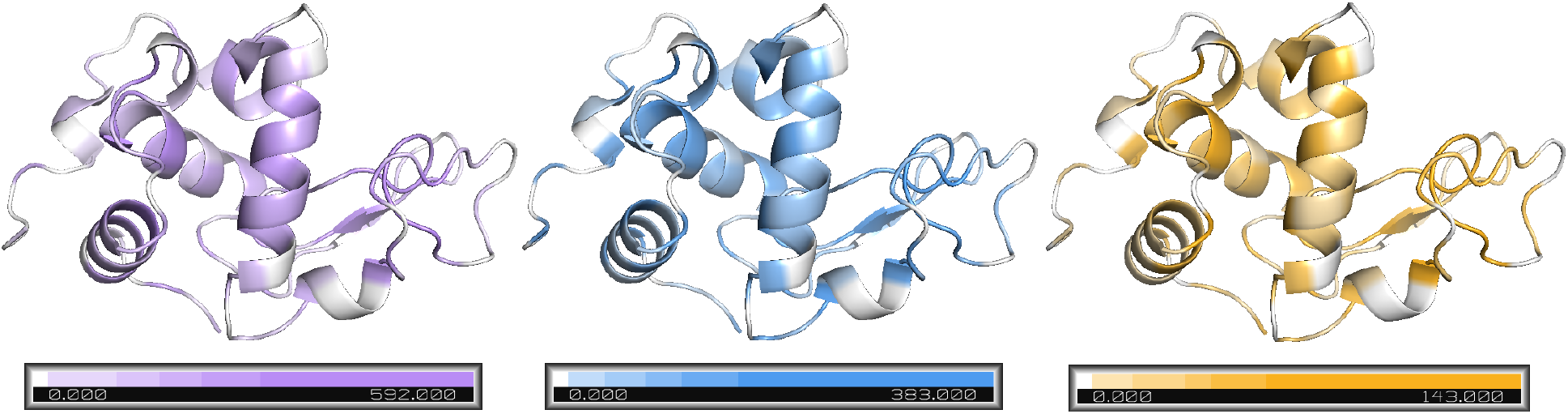
Per-residue topological participation in the Series (S), Parallel (P) and Cross (X) relation types, mapped onto the three-dimensional structure of hen egg-white lysozyme (PDB 1AKI, chain A). For each contact pair, the number of other contact pairs standing in the selected relation to it is counted, and that count is added to both of its endpoint residues. The value shown for a residue is the sum over the contact pairs it participates in. It is therefore a measure of topological participation and not a count of that residue’s own contacts. The 3D relation visualization currently supports only the three basic relation types; their concerted counterparts (CP, CP^−1^, CS) are not included. The Parallel mapping pools Parallel and Inverse Parallel relations. Residues with a value of zero are white. Each map is scaled to its own distribution; the colour bar in the viewer is labelled with 0 and the maximum of the scale, and the five level bounds are printed to the PyMOL log (S: 99, 161, 221, 306 and 592; P: 34, 72, 105, 158 and 383; X: 14, 32, 41, 60 and 143). Hence, identical colours in different panels do not represent identical values. Colouring is applied to a copy of the object, so the original structure and its deposited B factors are unchanged.

### 3.3 Multi-File Analysis: Tracking Topological Dynamics in Trajectories

Multi-file analysis extends circuit topology beyond a single structure: it batch-processes a folder of structures, or the frames of a molecular dynamics trajectory, in a single pass. We demonstrate the statistics output on hen egg-white lysozyme and the time-resolved output on a molecular dynamics trajectory of an intrinsically disordered protein undergoing compaction.

Multi-file analysis accepts either a folder of structures or a trajectory loaded into PyMOL, and we demonstrate both. The trajectory was loaded as a PDB file paired with its XTC trajectory file, while the hen egg-white lysozyme was loaded as part of a folder on a separate run. No conversion of the trajectory to individual PDB frames is required, as the plugin analyses the frames of the loaded object directly. For consistency, contact-pair parameters were configured with a 4.5 Å cutoff, five minimum atom-atom contacts, and three excluded neighbours. Upon executing the analysis with the ‘Statistics Plot’ and ‘P,S,X relations plotted over time’ options selected (to enable the latter, users must first check ‘P,S,X relation counts (.csv)’ in the export options), the plugin generated the outputs shown in Figs. 6 and 7.

**Fig. 6:**
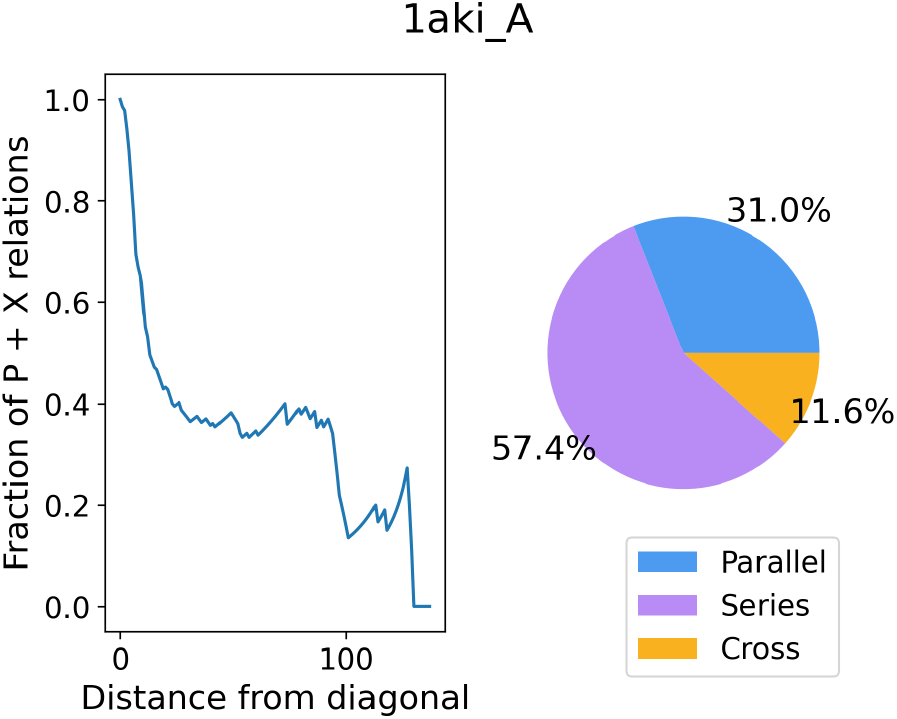
Statistics plot produced by multi-file analysis, shown here for hen egg-white lysozyme (PDB 1AKI, chain A). Left: the fraction of P + X relations, i.e. the share of contact-pair relations that are Parallel or Cross (non-Series), as a function of distance from the diagonal of the relation matrix; ProteinCT [6] calls this quantity the “entangled” fraction, a statement about relation types and not about geometric entanglement. Right: the distribution of relation types. Relation-type proportions follow the reference ProteinCT definition, in which the Parallel total also includes the concerted parallel types (CP, CP^−1^) and the Series total includes concerted series (CS). The per-residue maps of Fig. 5 count the plain relation types only.

**Fig. 7:**
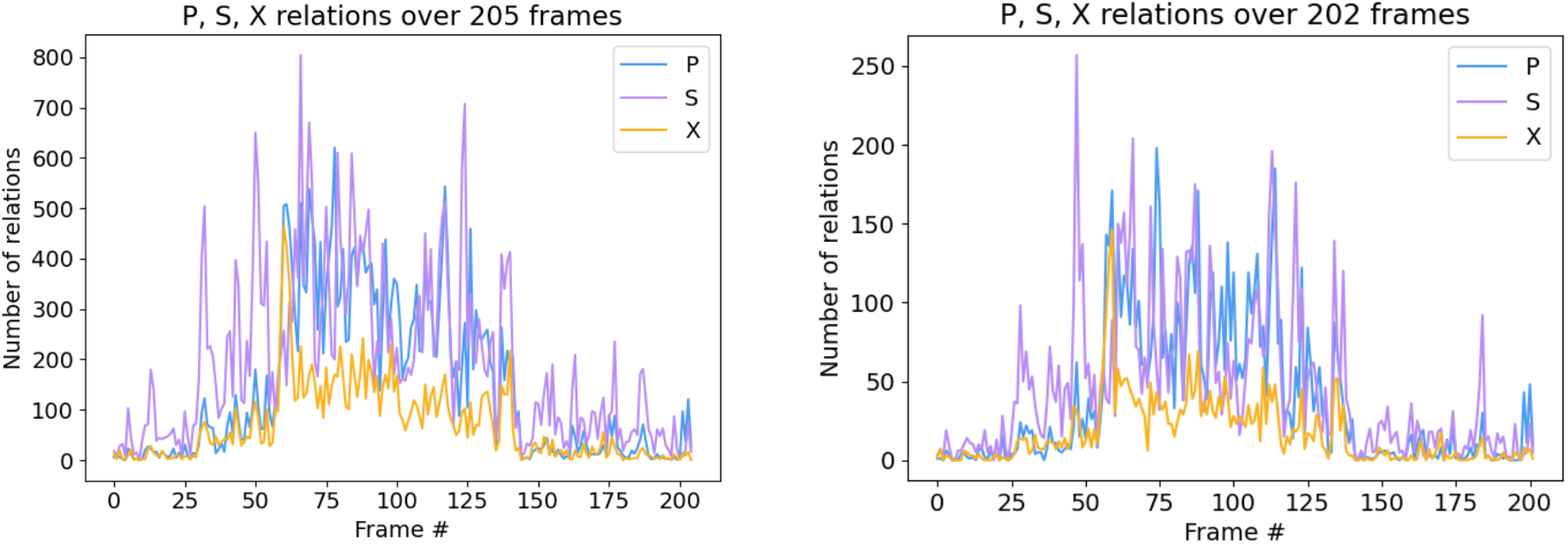
The number of Parallel (P), Series (S) and Cross (X) relations plotted over the 205 frames of the molecular dynamics trajectory of replica 3 of the AF1c S203P fragment. Left: Hydrogens included (by toggling on ‘Count hydrogen atoms (ProteinCT-identical)’ in the plugin GUI). Right: with the default heavy-atom criterion, three of the 205 frames contain no residue pair that satisfies the contact criterion. The plugin reports such frames and skips them, so only 202 frames reach the final time-resolved output.

The statistics plot (Fig. 6) summarises a structure in two ways. The fraction of P + X relations is plotted against the distance from the diagonal of the relation matrix, i.e. the separation of two contact pairs in the sorted contact list; it is highest for neighbouring contact pairs and falls with increasing separation, and its first and last points are fixed at 1 and 0 by construction. The relation-type distribution for lysozyme is 57.4% Series, 31.0% Parallel and 11.6% Cross.

Applied to replica 3 of the AF1c S203P fragment trajectory, the time-series output shows how the numbers of Series (S), Parallel (P) and Cross (X) relations evolve across the 205 simulation frames (Fig. 7). The number of relations of each type fluctuates throughout, which is consistent with the AF1c S203P fragment being an intrinsically disordered protein. Moreover, it can be observed in both plots (including and excluding hydrogen atoms) that the number of relations of all three types increases before returning to baseline, which would be consistent with a transient compaction of the AF1c S203P fragment. This example illustrates how multi-file analysis extends circuit topology from a static structural descriptor to a time-resolved probe of conformational change.

## 4. Impact

The Circuit Topology PyMOL plugin significantly lowers the technical barrier to applying protein CT analysis, thereby broadening the range of research questions that can be practically addressed. By embedding CT analysis within a premier molecular visualization platform, the plugin transforms specialized, script-based workflows into interactive, visually guided explorations of protein folding and functional regulation.

The plugin empowers researchers to perform residue- and region-specific investigations of topological motifs, allowing for deeper insights into the contributions of circuit topology to protein compaction and folding. Its support for trajectory-based analysis enables the systematic study of how topological relationships evolve over time in molecular dynamics simulations, particularly for intrinsically disordered proteins. Furthermore, the integration of the CT Folding Score provides a critical quantitative link between topological organization and molecular compaction, directly supporting emerging research into disorder–function relationships.

Technically, the plugin overcomes the limitations of previous methodologies that required substantial expertise in Python-based environments. It improves the pursuit of established research questions by providing an automated graphical interface that reduces setup time and minimizes data handling errors. This integration enables rapid iteration between structural inspection and topological quantification, making advanced descriptors accessible for routine use.

By making circuit topology analysis accessible within standard PyMOL workflows, the plugin transforms CT from a specialized post-processing step into an interactive component of everyday structural analysis. Users can readily visualize, quantify, and export topological features alongside conventional structural representations. This accessibility encourages a more frequent and exploratory use of CT descriptors across the structural biology community.

Finally, the plugin targets the large community of PyMOL users and is distributed openly with cross-platform support. Its reliance on established, citable CT methodologies facilitates seamless integration into publishable research. While designed for academic exploration, the tool may also be useful in applied settings such as drug discovery and protein engineering, where topological descriptors are used alongside conventional structural analysis.

## 5. Conclusions

In this work, we present a PyMOL plugin that makes protein CT analysis broadly accessible through a graphical, integrated, and automated workflow. By combining established CT methodologies with PyMOL’s ecosystem and visualization capabilities, the plugin enables intuitive exploration, quantitative analysis, and direct visual interpretation of protein topological organization across single structures, local regions, and molecular dynamics trajectories.

Validation against the reference ProteinCT implementation demonstrates exact agreement on the structures tested, and the lysozyme analyses reported here are reproducible from a versioned release with a single command (the trajectory analysis from the deposited AF1c dataset with the same release).

While the plugin substantially improves accessibility and usability, certain limitations remain. The plugin analyses the polymer component of protein structures supplied as PDB or mmCIF files, together with trajectories in the XTC format. Waters, ions, ligands and other non-polymer atoms are excluded automatically and reported. Nucleic acids and unknown residues are removed when the chain is parsed, so protein-nucleic acid complexes are analysed as protein only. Chemically modified residues deposited as heteroatom records, such as selenomethionine and phosphorylated residues, are removed with the ligands even though they occupy backbone positions, leaving a gap at that position. Contact pairs are detected between heavy atoms by default, so models with and without hydrogens give the same contact pairs; the optional all-atom mode reproduces the reference ProteinCT implementation on hydrogen-bearing files but is only comparable between files with the same hydrogen representation. Alternate conformations are taken as presented by the structure parser.

The plugin cannot analyze extremely small structures: at least two intra-chain contact pairs are required before any relation exists. The contact-length and contact-energy filters operate on single-chain structures only and are available during multi-file analysis only, aligned with the ProteinCT implementation. The energy filter additionally recognises only the twenty standard amino acids. Per-residue colouring and the CT Folding Score are computed for each chain separately, so relations between chains contribute to neither. Single-chain and multi-chain analyses use two different, non-interchangeable relation vocabularies (Section 2.1): inter-chain contact pairs are included in whole-model analysis, and the appropriate level is selected automatically from the number of chains present.

The reported quantities should be read as what they are. A contact pair is a threshold decision - a minimum number of atom-atom contacts within a distance cutoff. It is not a measurement of interaction strength. The per-residue values mapped onto the structure are sums over the relations in which a residue’s contact pairs participate. They are not counts of that residue’s own contacts. Each three-dimensional map is scaled to its own distribution, so identical colours in different maps do not represent identical values.

Analysis of a single small structure completes in well under a tenth of a second and uses approximately 43 MB of memory, essentially independently of workload. This scales with trajectory length: colouring an entire trajectory by relation type holds every frame in memory simultaneously and reaches approximately 1.3 GB for two hundred frames of a small protein. Therefore, very long trajectories should be coloured in segments. The plugin targets PyMOL 3.1.6.1 and installs its dependencies into PyMOL’s own environment. If PyMOL was installed system-wide this requires elevated admin privileges.

## Notes

### Competing Interest Statement

The authors have declared no competing interest.

### Summary of Updates

We applied minor changes to improve the article: - Improved terminology, consistency, and clarity throughout the manuscript and software. - Clarified methods and strengthened reproducibility and documentation.

