## Supplemental Material for "PyMOL plugin for Protein Circuit Topology"

**Supplementary Material**
***PyMOL plugin for Protein Circuit Topology***

Matīss Dimiņš, Alexander Bazba, Ádám Mogyorósi, Ella Kennon, Tomás Díaz Fiol, Leïla Aïkili Hagen, Vahid Sheikhhassani, Vasily Akulov, Alireza Mashaghi*

Faculty of Science, Leiden University, 2333 CC Leiden, The Netherlands

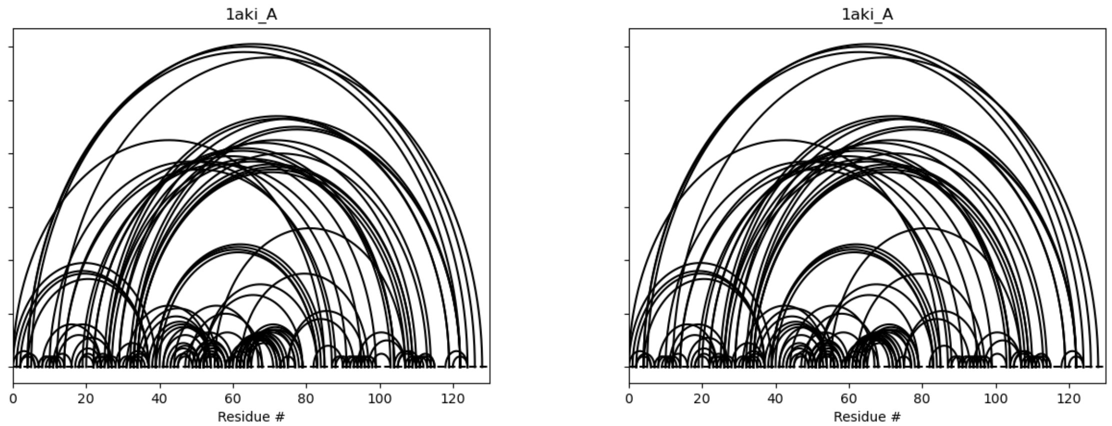


**Fig. S1**: Parity comparison between PyMOL ProteinCT plugin (left) and original ProteinCT method implementation (right) of circuit plots for hen egg-white lysozyme (PDB 1AKI).

| | [0 - 6] | [0 - 39] | [0 - 85] | [1 - 37] | [2 - 6] | [2 - 7] |
|---|---|---|---|---|---|---|
| [0 - 6] | - | CP | CP | X | CP-1 | X |
| [0 - 39] | CP-1 | - | CP | P-1 | P-1 | P-1 |
| [0 - 85] | CP-1 | CP-1 | - | P-1 | P-1 | P-1 |
| [1 - 37] | X | P | P | - | P-1 | P-1 |
| [2 - 6] | CP | P | P | P | - | CP |
| [2 - 7] | X | P | P | P | CP-1 | - |
| [2 - 37] | X | P | P | CP | CP-1 | CP-1 |
| [4 - 37] | X | P | P | CP | X | X |
| [4 - 122] | X | X | X | X | X | X |
| [4 - 124] | X | X | X | X | X | X |
| [5 - 126] | X | X | X | X | X | X |
| [7 - 11] | S | P | P | P | S | CS |
| [8 - 12] | S | P | P | P | S | S |
| [8 - 24] | S | P | P | P | S | S |
| [9 - 13] | S | P | P | P | S | S |
| [10 - 14] | S | P | P | P | S | S |

**Fig. S2**: Relation matrix export for hen egg-white lysozyme (PDB 1AKI, chain A), generated in .csv format for downstream statistical use. Contact-pair maps are arrays of 0's and 1's, which do not provide visual insight, so we have deliberately excluded them here.
